# Sensorimotor processing in the basal ganglia leads to transient beta oscillations during behavior

**DOI:** 10.1101/136879

**Authors:** Amin Mirzaei, Arvind Kumar, Daniel Leventhal, Nicolas Mallet, Ad Aertsen, Joshua Berke, Robert Schmidt

## Abstract

Brief epochs of beta oscillations have been implicated in sensorimotor control in the basal ganglia of task-performing healthy animals. However, which neural processes underlie their generation and how they are affected by sensorimotor processing remains unclear. To determine the mechanisms underlying transient beta oscillations in the local field potential (LFP), we combined computational modeling of the subthalamo-pallidal network for the generation of beta oscillations with realistic stimulation patterns derived from single unit data. The single unit data were recorded from different basal ganglia subregions in rats performing a cued choice task. In the recordings we found distinct firing patterns in the striatum, globus pallidus and subthalamic nucleus related to sensory and motor events during the behavioral task. Using these firing patterns to generate realistic inputs to our network model lead to transient beta oscillations with the same time course as the rat LFP data. In addition, our model can account for further non-intuitive aspects of beta modulation, including beta phase resets following sensory cues and correlations with reaction time. Overall, our model can explain how the combination of temporally regulated sensory responses of the subthalamic nucleus, ramping activity of the subthalamic nucleus, and movement-related activity of the globus pallidus, leads to transient beta oscillations during behavior.

**Significance Statement:** Transient beta oscillations emerge in the normal functioning cortico-basal ganglia loop during behavior. In this work we employ a unique approach connecting a computational model closely with experimental data. In this way we achieve a simulation environment for our model that mimics natural input patterns in awake behaving animals. Using this approach we demonstrate that a computational model for beta oscillations in Parkinson’s disease can also account for complex patterns of transient beta oscillations in healthy animals. Therefore, we propose that transient beta oscillations in healthy animals share the same mechanism with pathological beta oscillations in Parkinson’s disease. This important result connects functional and pathological roles of beta oscillations in the basal ganglia.

## Introduction

Exaggerated cortico-basal ganglia oscillations in the beta band (15 to 30 Hz) are a common feature of Parkinson’s disease (PD; Brown et al., 2001; Hammond et al., 2007; Levy et al., 2002). However, beta oscillations are not always pathological. Brief epochs of beta oscillations have been implicated in sensorimotor control in the healthy basal ganglia (Berke et al., 2004; Leventhal et al., 2012; Courtemanche et al., 2003; Feingold et al., 2015). These studies suggest that temporally regulated transient beta oscillations are important for normal functioning of the motor system.

The origin of beta oscillations in the cortico-basal ganglia system remains unknown. However, interactions between subthalamic nucleus (STN) and globus pallidus externa (GPe) can generate beta oscillations as has been shown in experimental (Bevan et al., 2002; Tachibana et al., 2011) and computational (Terman et al., 2002; Kumar et al., 2011, Pavlides et al., 2015; Wei et al., 2015) studies. Anatomically, STN and GPe are densely and reciprocally inter-connected (Shink et al., 1996). STN cells excite neurons in GPe (Kitai and Kita, 1987), which in turn receive inhibitory input from GPe (Smith et al., 1990; Parent and Hazrati, 1995). Such recurrent excitation-inhibition can generate oscillations (Plenz and Kitai, 1999; Brunel, 2000), which may then propagate to other regions in the cortico-basal ganglia loop.

Beta oscillations have been proposed to play a functional role in maintaining the status quo in the motor system (Engel and Fries, 2010; Gillbertson et al., 2005). This idea has been supported by increased cortical beta-band activity during maintenance of a static position (Baker et al., 1997), active suppression of movement initiation (Swann et al., 2009), and post-movement hold periods (Pfurtscheller et al., 1996). Accordingly, beta power decreases in the cortico-basal ganglia loop during movement preparation and execution (Sochurkova and Rektor, 2003; Pfurtscheller et al., 2003; Alegre et al., 2005; Kuhn et al., 2004). However, recent studies have indicated a more complex picture in which beta oscillations affect behavior through motor adaptation (Tan et al., 2014) and modulation of task performance (Feingold et al., 2015).

Supporting a more complex picture of beta oscillations, we provided evidence that basal ganglia beta oscillations are involved in sensorimotor processing and the utilization of cues for behavior (Leventhal et al., 2012). In particular, we found that beta power increases following sensory cues and movement initiation depended on how fast the animals reacted to a sensory cue. For short reaction times, LFP beta emerged after movement initiation, whereas for long reaction times, two separate beta epochs occurred, one before and one after movement initiation. In addition to modulation of beta power, we also observed that beta phases were affected by task events differently. Sensory cues, but not movement initiation, lead to a short-latency phase reset in the beta band (Leventhal et al., 2012).

These complex oscillatory dynamics present both a challenge, and an opportunity, for understanding underlying cortico-basal ganglia circuit mechanisms. Currently, it is unknown whether pathological beta oscillations in Parkinson’s disease share the same mechanisms with transient beta oscillations in healthy animals. If this is the case, computational models for beta oscillations should be able to account for the complex beta dynamics in both healthy and Parkinsonian animals. Recent network models of beta oscillations in Parkinson’s disease have emphasized that besides structural changes (e.g. connection strengths), changes in spiking activity of external inputs can promote beta oscillations (Kumar et al., 2011), which might drive transient beta oscillations. Here we exploit this property by directly using activity patterns recorded in healthy rats during task performance (Schmidt et al., 2013; Mallet et al., 2016) as input to our computational model to study the resulting impact on the beta dynamics. Employing this novel approach we find that our model can account for the complex beta dynamics in the healthy rat LFP. Our results support overlapping mechanisms for pathological and healthy beta oscillations and provide the basis for studying the functional role of beta oscillations in network models.

## Materials and methods

### Network model

The basic model structure and the parameter settings are the same as in Kumar et al. (2011). Briefly, the model includes 1000 excitatory STN neurons, and 2000 inhibitory GPe neurons. Neurons were implemented as leaky integrate-and-fire neurons. Synaptic input was modeled as transient exponential conductance changes. All model neurons receive uncorrelated Poisson spike trains as inputs so as to achieve previously reported baseline activities for STN (15 Hz) and for GPe (45 Hz; Bergman et al., 1994; Raz et al., 2000). All network simulations were written in python using pyNN as an interface to the simulation environment NEST (Gewaltig and Diesmann, 2007). Analysis of the simulation results and the LFP and single unit data were performed using MATLAB R2013b (version 8.2.0.701; The MathWorks Inc., Natick, MA).

For the model variant without recurrent connections in STN (Figure 8), we used slightly different parameters for the connection probabilities, synaptic weights and transmission delays (Table 1). Furthermore, the background Poisson input to the model neurons was adjusted so that the neurons had a broader distribution of baseline firing rates that closer matched the firing rate distribution in the rat data (Schmidt et al., 2013; Mallet et al., 2016).

**Table 1:** Comparison of model parameters in Kumar et al. (2011) and the modified model without recurrent STN connections.

| Kumar et al., 2011 | Modified model |
| --- | --- |
| $CP_{STN-STN} = 0.02$ | $CP_{STN-STN} = 0$ |
| $CP_{STN-GPe} = 0.02$ | $CP_{STN-GPe} = 0.022$ |
| $CP_{GPe-STN} = 0.02$ | $CP_{GPe-STN} = 0.035$ |
| $CP_{GPe-GPe} = 0.02$ | $Cp_{GPe-GPe} = 0.02$ |
| $J_{STN-STN} = 1.2$ | $J_{STN-STN} = -$ |
| $J_{STN-GPe} = 1.2$ | $J_{STN-GPe} = 1.2$ |
| $J_{GPe-STN} = -1.135$ | $J_{GPe-STN} = -0.8$ |
| $J_{GPe-GPe} = -0.725$ | $J_{GPe-GPe} = -0.725$ |
| $d_{STN-STN} = 2$ | $d_{STN-STN} = -$ |
| $d_{STN-GPe} = 5$ | $d_{STN-GPe} = 6$ |
| $d_{GPe-STN} = 5$ | $d_{GPe-STN} = 6$ |
| $d_{GPe-GPe} = 2$ | $d_{GPe-GPe} = 3$ |

CP: Connection Probability, J: Synaptic weight, d: Delay (in ms)

### Experimental design and statistical analysis

We combined previously recorded data sets of tetrode recordings in different basal ganglia subregions of rats performing a stop-signal task (for details see Leventhal et al., 2012; Schmidt et al., 2013; Mallet et al., 2016). To exclude potential multi-unit activity from our recordings, we only included units with less than 1% of inter spike intervals shorter than 1 ms in our data set. The combined data set contained 226 STN units from overall 40 recording sessions in 5 different rats, 149 putative prototypical GPe units from 41 recording sessions in 4 different rats, and 326 putative MSNs from 97 recording sessions in 9 different rats. Between two recording sessions tetrodes were typically moved by at least 80*μ*m, and we therefore considered units recorded in different sessions as different units. Animals performed a stop-signal task, but here we only analyzed the subset of correct Go trials in which the animal moved contralateral to the recording site.

To identify STN neurons responding to the Go cue instructing contralateral movement (Figures 1C, D), we used a shuffle test to determine whether neural activity significantly increased within 150 ms after the Go cue. The time of each spike within -500 ms to +200 ms relative to the Go cue was changed to a random spike time within the same time window. Then we compared the number of actual spikes with the number of shuffled spikes in small time windows after the Go cue (15 non-overlapping 10 ms windows from 0 to 150 ms after the Go cue). We repeated this procedure 10000 times and used the fraction of shuffles in which the number of shuffled spikes exceeded the number of actual spikes as the p-value to estimate statistical significance. STN neurons showing a p-value less than 0.05/15 for at least one bin after the time of the Go cue were considered sensory responsive. We performed the same shuffling method on GPe neurons to select movement responsive GPe neurons (Figure 1F), using all spikes within -1s to +1s relative to movement onset to detect firing rate changes for 50 ms time windows from 0 to 250 ms after movement onset (i.e. 5 non-overlapping time bins). GPe neurons showing a p-value less than 0.05/5 for at least one bin after movement onset were considered as movement responsive.

**Figure 1:**
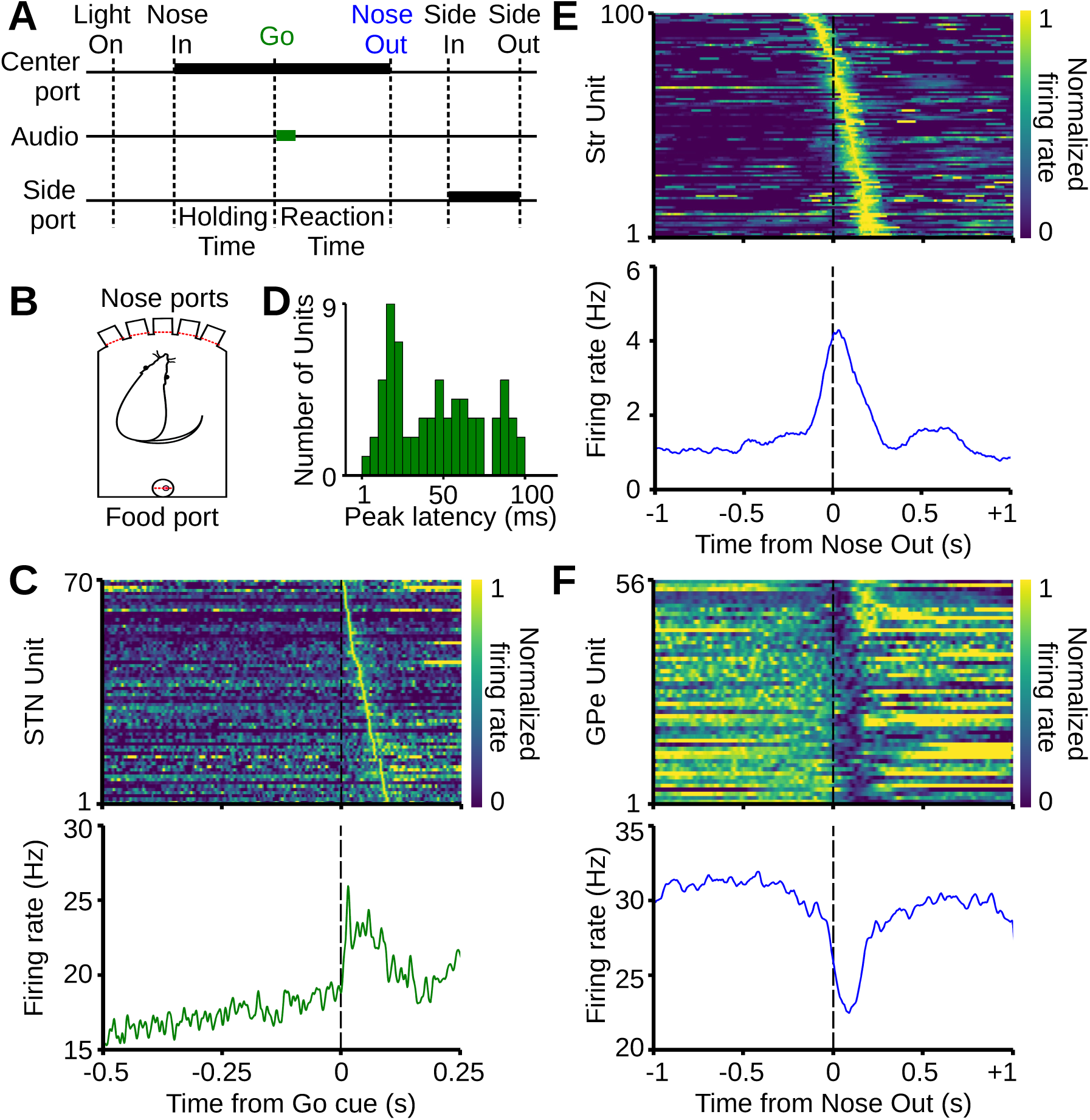
Single unit responses to sensory and motor events during performance of the behavioral task. **A**, Sequence of behavioral events during the experiment. Thick black bars show the position of the animal and thick green bar shows the occurrence of the sensory cue. Holding time refers to a random time delay (500 to 1200 ms) in which the animal waits in one of the three central ports for the sensory cue. Reaction time is measured as the time between the onset of the Go cue and movement initiation (Nose Out). **B,** Scheme of the operant chamber with five nose ports in front and a food port in the back. **C** (top) Normalized mean firing rates of single STN units responding to the Go cue with an increase in firing rate (sorted by peak latency; each row shows activity of one unit). Bottom, corresponding mean firing rate of the STN subpopulation. **D**, Distribution of peak latencies relative to the time of Go cue for STN neurons shown in C. **E** (top) Normalized firing rates of single units in the striatum (putative MSNs) increasing their activity around movement onset (sorted by time of peak activity). Bottom, corresponding mean firing rate of the subpopulation. **F**, Same as E, for GPe subpopulation decreasing activity around movement onset.

To identify movement-responsive MSNs in our single unit data, average firing rates of MSNs were sorted based on their peak time within the interval from one second before to one second after movement initiation. MSNs with a peak firing rate between 150 ms before to 150 ms after movement onset were considered as movement-responsive MSNs (n = 100; see Figure 1E).

To determine whether a recorded unit showed a ramping firing pattern, we computed the average firing rates of each unit from one subregion over trials with a 50 ms sliding time window moving in steps of 10 ms from 1 s before the time of Go cue to the time of Go cue. Each resulting average firing rate was then normalized to values between 0 and 1 and then mean-subtracted before applying principal component analysis. First, we computed the corresponding covariance matrix of all normalized zero-mean firing rates. and then performed eigendecomposition on the covariance matrix using the *eig* function of MATLAB. The projection *p* of each normalized zero-mean average firing rate *r* to the first eigenvector (corresponding to the maximum eigenvalue) was then computed as the normalized dot product: *p*_*i*_*=⟨r*_*i*_,*v*_1_⟩/λ_1_ where *i* is the unit index and *v*_1_ the eigenvector with the largest eigenvalue*λ*_1_. This yielded one projection value *p*_*i*_ for each recorded unit. As the first eigenvector had a positive ramp over time, positive and negative projection values corresponded to positive and negative activity ramps of a recorded unit over time, respectively. The standard deviation of the projection distribution from a random covariance matrix is 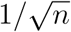 (Anderson, 2003), with *n* being the number of units. We considered neurons with a projection larger than 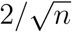 or smaller than 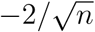 as positive and negative ramp neurons, respectively (Figures 2A, B). This analysis method was applied to determine positive and negative ramps in GPe and STN.

**Figure 2:**
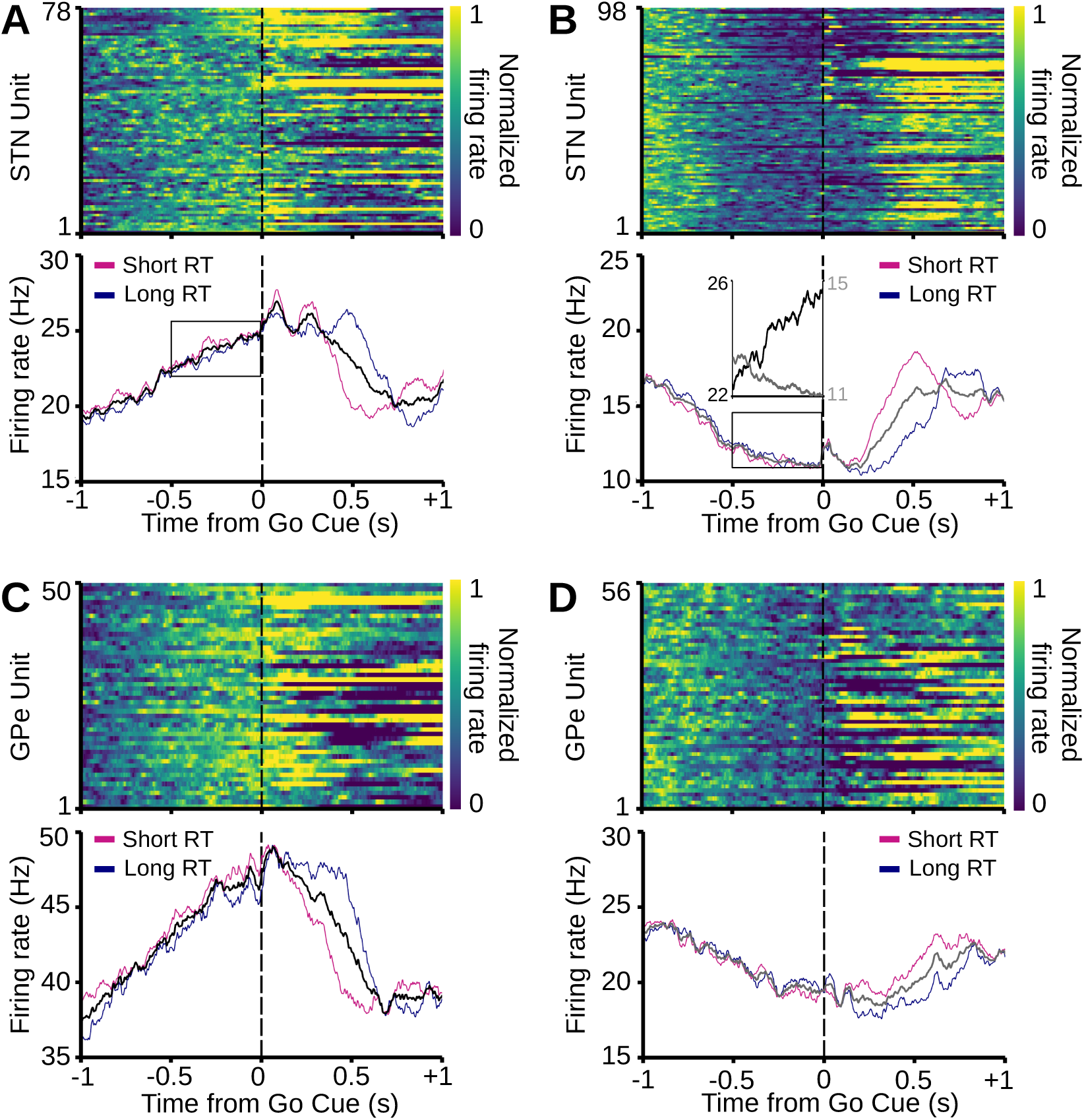
Ramping activity in STN and GPe while the animal is waiting for the Go cue. **A** (top) Normalized mean firing rate of single STN units with a positive ramp in firing rate before the Go cue. Bottom, corresponding mean firing rate of the STN subpopulation in all trials (black) and subsets of long (cyan) and short (magenta) reaction time trials. **B**, Same as A, for single STN units with a negative ramp in their firing rate before the Go cue. Inset, direct comparison between average firing rates of neurons in A, and B, corresponding to the areas inside the black rectangles. **C**, **D**, Similar to A and B, respectively, for GPe units.

### Modeling of sensory responses

To simulate sensory responses of STN neurons to the Go cue (Figures 1C, D), we used inhomogeneous Poisson generators, each of which targeted one STN neuron in the model. The firing rate modulation of each inhomogeneous Poisson generator was a half sine wave with a duration of 20 ms and maximum amplitude of 180 Hz. The latency of the sensory stimulation for each STN neuron in the model was considered as the time interval between the peak of the half sine wave and the time of the Go cue, which was taken randomly from the latency distribution of the sensory STN neurons in our experimental data (Figure 1D). Since in our single unit data 30% of the STN neurons responded to the Go cue, for each simulation we targeted 30% of randomly chosen STN neurons (as “sensory” STN neurons) in the network model. Thereby, sensory responses in STN neurons could propagate in our network model to GPe, similar to some short latency responses we previously reported in GPe (Schmidt et al.,2013).

### Modeling of motor responses

Firing rates of the movement-responsive MSNs (Figure 1E) were summed up and used as the firing rate pattern of an inhomogeneous Poisson generator representing striato-pallidal movement-related inhibition in the network model. Since 38% of the GPe neurons in our experimental data showed movement-related inhibition (Figure 1F), for each simulation we targeted a randomly chosen 38% of the GPe neurons (as “motor” GPe neurons) in the network model.

### Modeling of firing rate ramps

To simulate the positive and negative ramps in the activity of the STN neurons observed before the Go cue (Figures 2A, B), for each simulation, we divided STN neurons in the network model into two non-overlapping subpopulations. The fraction of STN neurons in each subpopulation in the network model was similar to the fraction we obtained from our experimental data (i.e. 34% of neurons exhibited a positive ramp, 43% a negative ramp). We used an inhomogeneous Poisson generator with a positive ramp firing rate pattern as excitatory input to the positive ramp STN subpopulation in the model. The positive ramp in the firing rate of the inhomogeneous Poisson generator started 500 ms before the Go cue at 0 Hz and reached 250 Hz at the time of the Go cue and stayed constant until the movement onset (Figure 3B). Such a stimulation lead to a 4 Hz increase in the activity of the positive ramp STN subpopulation in the network model during the 500 ms time interval preceding the Go cue, similar to what we observed in our experimental data (Figure 2A).

**Figure 3:**
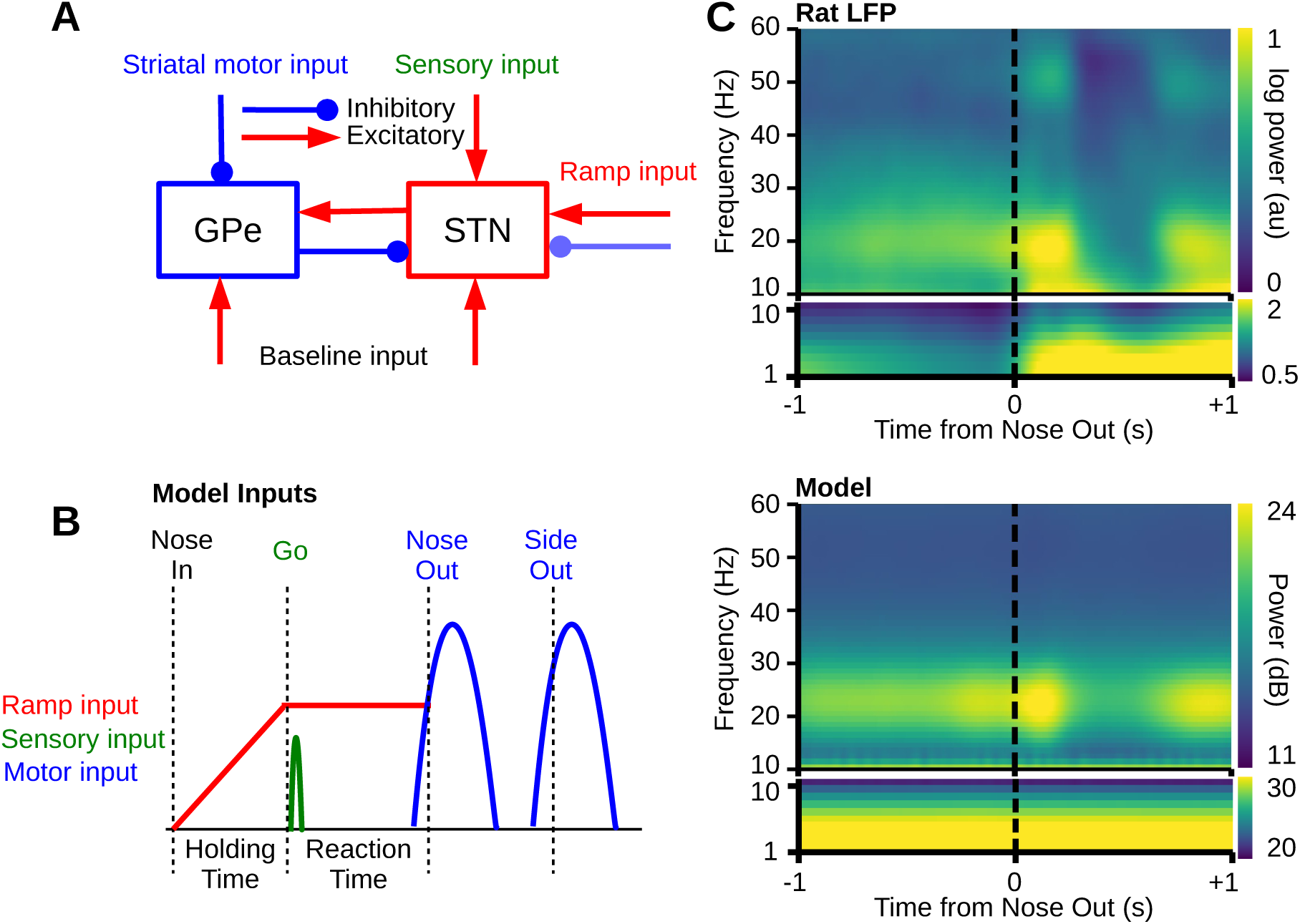
Computational model of beta oscillations stimulated with biologically realistic input patterns. **A**, Scheme of the STN-GPe spiking neuronal network model. Motor input is provided as striatal inhibitory input to the GPe whereas sensory input is provided as excitatory input to the STN. Ramp input comprises separate excitatory and inhibitory inputs to separate STN subpopulations (see Methods). **B**, Schematized temporal sequence of inputs to the network model during simulation of the behavioral task. **C** (top) Mean spectrogram of GPe LFP data showing modulation of LFP beta power during movement initiation. Bottom, Mean spectrogram (over 400 simulations) of GPe average firing rates for simulation of correct Go trials in the network model matching the time course of beta power in the experimental data.

Similarly, to simulate the negative ramp in the activity of STN neurons, we used another inhomogeneous Poisson generator with a positive ramp firing rate pattern as inhibitory input to the negative ramp STN model neuron subpopulation. The positive ramp in the firing rate of the inhibitory inhomogeneous Poisson generator started 500 ms before the time of Go cue at 0 Hz and reached 350 Hz at the time of the Go cue and stayed constant until the movement onset. Such a stimulation pattern lead to a 1 Hz decrease in the activity of the negative ramp STN neurons in the network model during 500 ms time interval preceding the Go cue, similar to what we observed in our experimental data (Figure 2B).

### Time-frequency analysis

The power spectrogram was computed by convolving 10 seconds of the GPe population firing rate (from -5 to +5 seconds relative to the time of movement onset) in the model with a standard Morlet wavelet (*σ* = 0.849*/f*) of integer frequencies (*f* = 1 *to* 500 *Hz*), and taking the logarithm of the squared magnitude of the resulting time series. To generate Figure 3C, bottom, we computed the mean spectrogram across 400 simulations of the model. The same method was used for GPe LFP data to generate Figure 3C, top. For each time point in the spectrogram, we summed the power in the beta range (15 to 30 Hz) and divided it by the summed power across all frequencies (1 to 500 Hz) to obtain continuous relative beta power, shown in Figures 4A, 4B, 4E, 4F, and 6B.

**Figure 4:**
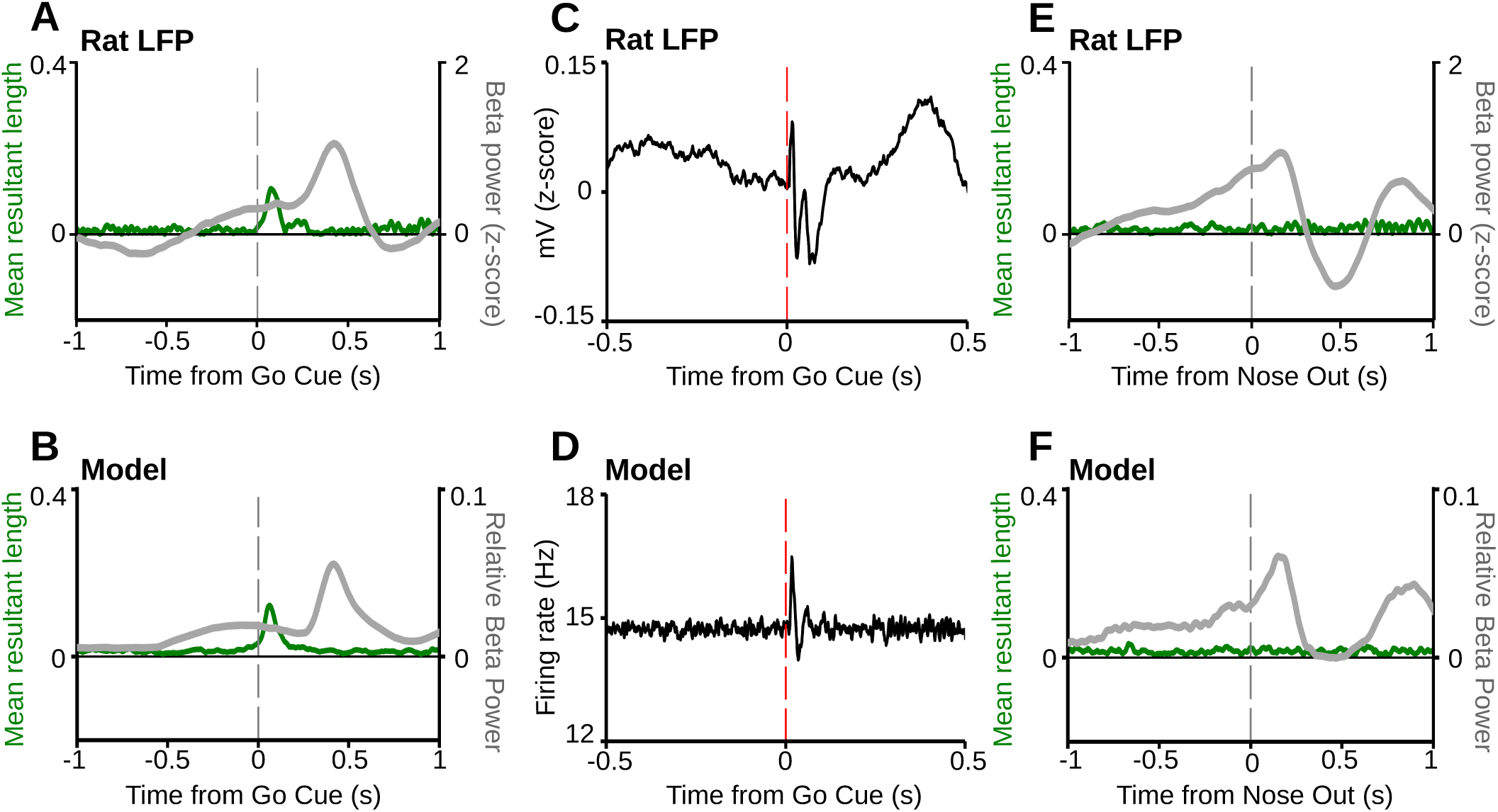
Sensory cues lead to a beta phase reset in both experimental data and in the network model. **A, B**, Time resolved beta mean resultant length (left axes, green) and beta power (right axes, gray) of GPe LFP data during correctly performed contralateral go trials averaged across all rats (A) and of the network model GPe population firing rate (B; average of 400 simulations). Note that sensory input is associated with a phase reset in both experimental data and in the model, shown as a brief increase in the value of the mean resultant length after the Go cue. **C**, Mean of the raw experimental STN LFP data, over all correctly performed contralateral go trials, aligned to the Go cue. D, Mean of the STN population firing rates in response to the Go cue in the network model (average of 400 simulations). **E, F**, The same analysis and simulations as in A and B, respectively, but aligned to movement onset. Note that the phase distribution is random during initiation and execution of movement in both the rat and the network model (no increase in the mean resultant length around movement onset).

### Mean resultant length

The GPe population firing rate in the network model was convolved with the standard Morlet wavelet of each integer frequency in the beta band (15 to 30 Hz). For each frequency, the Hilbert transform of the filtered signal was computed to obtain a phase over time. The phase spread for each time point was then calculated by computing the length of the mean resultant vector over all trials using 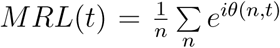, where *θ* (*n, t*) is the phase of the n*th* trial at time *t* (*n* = 400 for the model). This results in a continuous measure of phase spread for each frequency in the beta range. The mean resultant lengths shown in Figure 4 were computed by taking the average across all beta frequencies.

## Results

To determine whether a computational model for pathological beta oscillations in the STN-GPe network (Kumar et al., 2011) can account for complex beta dynamics during behavior in healthy animals, we devised realistic stimulation patterns for the network model based on single unit recordings in rats performing a cued choice task (Schmidt et al., 2013; Mallet et al., 2016). At the beginning of each trial, the rat entered one of three center nose ports in an operant chamber (“Nose-in” event; Figures 1A, B). The rat was trained to then hold its position for a variable time interval (“Holding time”; 500-1200 ms) until a Go cue instructed the rat to quickly move its head to the adjacent left or right side port (“Nose-out” event; Figures 1A, B). Correct performance of the task was rewarded with a sugar pellet. While the animals performed the task we recorded in the striatum, GPe and STN to determine activity patterns of single units during the time of the Go cue and during movement initiation. Then we used these activity patterns to construct realistic input patterns for our network model. The network model we use here is a large-scale spiking network model consisting STN and GPe populations with conductance based synapses (Kumar et al., 2011; see Methods). Stimulating the network model via the realistic stimulation patterns allowed us to compare the resulting oscillatory dynamics in the model with properties of oscillations in the rat LFPs.

### Brief, short-latency sensory responses in STN

30% (70/226) of STN units responded to the Go cue with an increase in firing rate (Figure 1C; shuffle test, p*<*0.05/15; see Methods). In line with our previous reports on a subset of the same data (Schmidt et al., 2013), this included units with a very short latency (around 10-30 ms), and responses of individual units were typically very brief (see Figure 1C, top panel). A potential source of such short latency sensory responses of the STN units is pedunculopontine tegmental nucleus (PPN; Pan and Hyland, 2005). In addition to the short latency responses of the STN units, some STN units responded with a longer latency (around 40-100 ms), so that the overall distribution of peak response latencies had a bimodal shape (Figure 1D). To mimic this STN response pattern to salient sensory stimuli, individual STN units received brief excitatory pulses with a fixed latency sampled from the latency distribution. These pulses were then used as input to 30% randomly chosen STN model neurons (“sensory” STN neurons) to match the fraction of responding STN units in our single unit data.

### Movement-related activity in striatum and GPe

30% (100/320) of putative medium spiny neurons (MSNs) in the striatum increased their activity during contralateral movements (Figure 1E; see Methods; also see Schmidt et al., 2013). We focused here on contralateral movements as most neurons typically responded more during contralateral than ipsilateral movements (Gage et al., 2010; Schmidt et al., 2013). In GPe, 38% (56/149) of the units decreased their activity during contralateral movements (Figure 1F; shuffle test, p*<*0.05/5; see Methods), possibly reflecting input from indirect pathway MSNs. Therefore, we assumed in the network model that striato-pallidal inhibition drives the GPe firing rate decreases during movement. We implemented this by generating inhomogeneous Poisson spike trains with a rate modulation following the MSN firing pattern during movement (Figure 1E). These spike trains were then used as inhibitory inputs to 38% of the network model GPe neurons (“motor” GPe neurons) to match the fraction of GPe units with movement-related firing rate decreases in the single unit data. Note that we restricted our analysis of GPe units to putative prototypical neurons (Mallet et al., 2016) because they receive input from MSNs and project to STN, while arkypallidal GPe neurons probably receive different inputs and do not project to STN (Mallet et al., 2012; Dodson et al., 2015).

### Ramping activity in STN and GPe while rats wait for the Go cue

In addition to single unit responses that could be classified as sensory or motor, in STN and GPe we found many units which exhibited a firing pattern that resembled a “ramp”, a continuous change in firing rate. A ramping pattern was present in the activity of 77% (176/226) of the STN units with either significantly increasing (positive ramp) or decreasing (negative ramp) firing rate while the animal was waiting for the Go cue (Figures 2A, B). Among the 176 ramping STN units, 44% (78/176) showed positive ramps (Figure 2A), whereas 55% (98/176) showed negative ramps (Figure 2B). However, the mean firing rate increase for the positive ramp units was four times as high as the mean firing rate decrease for the negative ramp units (4 Hz increase vs. 1 Hz decrease; inset in Figure 2B, bottom). The positive ramp was also observed in the average firing rate of the whole STN population starting 500 ms before the Go cue (data not shown). Functionally, these ramps may correspond to a brake signal, preventing premature movement initiation (Frank, 2006).

We found a similar pattern in the GPe with 71% (106/149) of the units exhibiting a significant ramping activity before the Go cue (Figures 2C, D). Among these, 47% (50/106) showed positive ramps (Figure 2C) and 52% showed negative ramps (Figure 2D). Similar to the STN units, on average, the amplitude of the positive ramp in GPe was four times as high as the amplitude of the negative ramp, resulting in a net positive ramp in the population activity (data not shown). One property of the positive ramp STN and GPe units was that in long reaction time trials their activity remained elevated after the Go cue (Figures 2A, C, bottom panels). This property played a key role for the beta dynamics in the model below.

Based on these ramping patterns in STN and GPe, we designed inputs to the model STN neurons that lead to similar activity ramps (see Methods). Due to the excitatory drive from STN to GPe, in the model the ramps in STN activity resulted in corresponding ramps in GPe.

### Sensorimotor model inputs modulate time course of beta oscillations

As a previous modeling study demonstrated that excitatory input to STN or inhibitory input to GPe can induce transient beta oscillations (Kumar et al., 2011), we hypothesized that the sequence of ramp, Go cue and movement-related activity patterns (Figures 3A, B) accounts for the complex beta dynamics in the LFP (Leventhal et al., 2012). First, we reproduced the time course of beta power modulation during movement initiation (Leventhal et al., 2012) using an extended data set of GPe recordings (Schmidt et al., 2013; Mallet et al., 2016). In the rat LFPs beta power started to increase before the time of movement initiation and then showed a pronounced peak just after movement onset (Figure 3C, top). The time course of beta power in the network model exposed to our single-unit stimulation patterns (Figure 3B) matched the experimentally observed results (Figure 3C, bottom), including the pre-movement beta power increase, the pronounced beta peak during movement, and the second beta peak related to the movement out of the side port (see Methods). The network model beta time course was in this case determined by the STN ramping activity, combined with the sensory responses of the STN neurons and the striato-pallidal motor inputs (Figure 3B). This is an important result because it connects single unit activity during task performance with oscillatory network dynamics.

Here we compared the experimental LFP data with the model population firing rate (Figure 3C). However, the origin of the LFP and its relation to spiking activity are not well understood in the basal ganglia. It seems that the LFP mostly reflects synchronized postsynaptic currents (Niedermeyer and Lopez da Silva, 1998; Nunez and Srinivasan, 2005; Jensen et al., 2005; McCarthy et al., 2011). However, we found that the time course of beta oscillations was very similar, irrespective of whether we used the population firing rate or the summation of inhibitory or excitatory postsynaptic currents to represent the experimental LFP data (data not shown). Therefore, to stay consistent with previous models (e.g. Kumar et al., 2011; Pavlides et al., 2015; Nevado-Holgado et al., 2014) we continue to use the population firing rate in the model to determine the presence of beta oscillations.

### Sensory responses in STN lead to a beta phase reset

In addition to the described changes in beta power, the phases of beta oscillations can be modulated by specific events in the behavioral task. Sensory cues (like the auditory Go cue) that did not lead to a distinctive increase in beta power were nevertheless followed by a short-latency phase reset in the LFP (Leventhal et al., 2012). By contrast, beta power increases during movement were not accompanied by a phase reset in the beta band (Leventhal et al., 2012). Here, we confirm this result for GPe recording sites using an extended data set (Figures 4A, E; Schmidt et al., 2013; Mallet et al., 2016). To determine which properties of the neural signal lead to a phase reset or to a power increase in the beta band, we calculated grand averages of raw LFP traces (Figure 4C). We found that briefly after the Go cue a single beta cycle was visible. This short oscillation was rather weak and could only be visible when looking at the mean of the LFP data over many trials (Figure 4C). This brief beta epoch was associated with beta phase reset in the LFP data, following the Go cue (Figure 4A). Interestingly, providing brief stimulation to the “sensory” STN neurons in the model leads to a brief low-amplitude beta oscillation, which also only became visible when inspecting the mean population firing rate over many stimulations (Figure 4D). Similar to the experimental data, “sensory” stimulation of the model STN leads to beta phase reset in the ongoing activity of the network model (Figure 4B). Therefore, we conclude that brief excitatory inputs to STN can induce weak and brief, phase-locked beta oscillations in the STN-GPe network, mimicking the experimentally observed results.

Beta elevation around the time of movement onset was not accompanied by a phase reset in both the rat LFP data and in the model (Figures 4E, F). It might seem counterintuitive that a strong stimulation leading to a clear increase in beta power did not reset the phase, whereas a weaker stimulation did. However, STN neuronal responses to the Go cue are brief, compared to the longer movement-related increases in the activity of MSNs (Figures 1C-E). Therefore, we hypothesized that the duration of neural responses to sensory and motor events might be the key difference. To test this, we systematically varied the duration of the inputs to the model “sensory” STN neurons and “motor” GPe neurons (note that the inputs are inhomogeneous Poisson spike trains with firing rate patterns of a half cosine wave; see Methods). We found that for brief inputs (leading to brief changes in the neuronal activity) there was a phase reset in the ongoing activity of the network model (Figure 5). Longer stimulations of “motor” GPe neurons elevated the beta power without phase reset (Figures 5C, D). For stimulation durations longer than a single beta period in the model (i.e. about 50 ms), we only observed beta power elevation without phase reset (Figures 5C, D). In fact, the maximal phase reset in the network model occurred when the stimulation duration was 25 ms, equaling half the beta cycle (Figures 5B, D). For the short stimulation duration the time to get to the maximum of the half cosine firing rate pattern is short (i.e. the slope is steeper). This effectively leads to no trial-to-trial variability because all realizations of the Poisson process with such a brief firing rate pattern are very similar (with respect to the spike times). This similarity in the input then leads to a similar response in the network model and therefore a phase reset across trials. In contrast, for longer stimulation the time to get to the maximum of the half cosine firing rate pattern is longer (with shallower slope). This leads to more trial-to-trial variability with respect to the spike times in the realization of the Poisson process. Correspondingly, this translates into trial-to-trial variability in the response of the network model to the long stimulation and therefore a random phase across trials.

**Figure 5:**
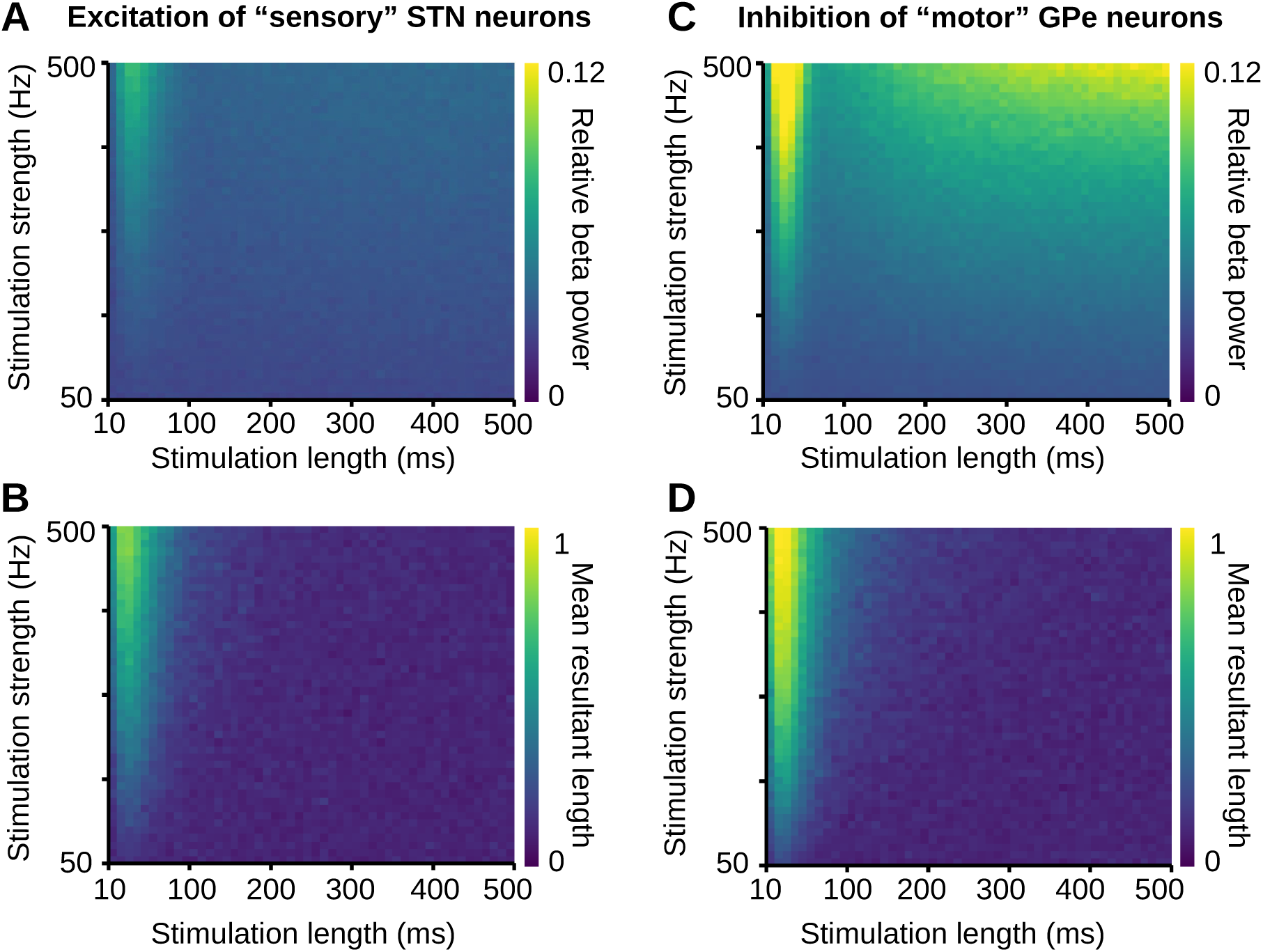
Effect of stimulation duration on beta power and phase reset in the network model. **A, B**, Relative beta power (A) and beta phase reset (B; measured by the mean resultant length) in the model GPe caused by excitatory input to the 30% “sensory” STN neurons of varying duration (x-axis) and strength (y-axis). **C, D**, Relative beta power (C) and phase reset (D) in the model GPe caused by inhibitory input to the 38% “motor” GPe neurons (see Methods) of varying duration (x-axis) and strength (y-axis). Note that in all panels we measure beta oscillations based on the GPe population firing rate.

Longer stimulations of the “sensory” STN neurons did not elevate the beta power in the network model (Figure 5A). This is because “sensory” STN units made up a smaller fraction (30%) of the STN population in our model compared to 38% “motor” GPe units (see above). The long stimulation of a small fraction of the STN neurons was not sufficient to bring the network model into the oscillatory state. In general, for a certain stimulation strength, the fraction of stimulated neurons in the network model is a key parameter determining the amount of evoked beta power (Kumar et al., 2011).

### Disentangling the complex relationship between reaction time and beta dynamics

The time course of beta oscillations depends on how fast the animal initiates movement in response to the Go cue (Leventhal et al., 2012). For short reaction times, the mean LFP beta power shows a single peak after movement initiation. For long reaction times, the mean LFP beta power shows two peaks, with the first peak before and the second peak after movement initiation (see highlighted 300 ms epochs preceding and following Nose Out in Figure 6A, right; see also Leventhal et al., 2012). The bimodal shape of the mean beta power for long reaction time trials is also visible when aligned to the Go cue (Figure 6A, left). A straightforward idea would be that the first peak of the mean beta power for long reaction time trials is mostly driven by the Go cue or, alternatively, by the upcoming movement. However, if the beta peak was driven by the Go cue, we would expect a higher peak for the data aligned to the Go cue than for the data aligned to movement onset. Accordingly, if the beta peak was related to the movement, we would instead expect a higher peak for the data aligned to the movement onset. In contrast, despite variability in reaction time, this peak had a similar shape and amplitude for both alignment to the Go cue and to movement onset. Therefore, this beta peak does not seem to be simply driven by a sensory or motor event. With the help of our network model, we disentangle the mechanisms underlying these reaction time-dependent complex features of beta.

**Figure 6:**
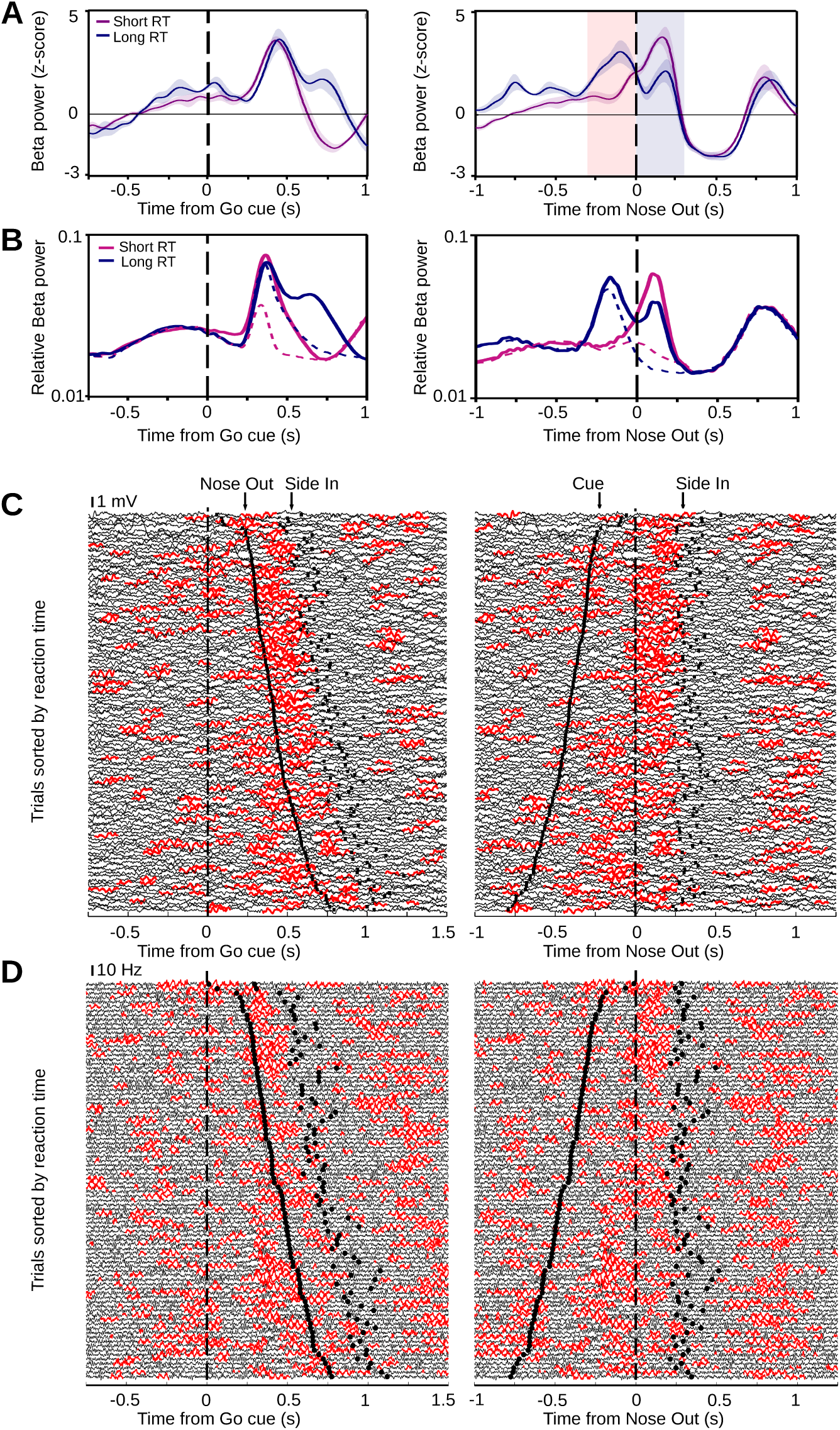
Relationship between beta oscillations and reaction time. **A**, Mean beta power of striatal LFP data for short (*<*500 ms) and long (*>*500 ms) reaction time trials aligned to the Go cue (left) and movement onset (right), averaged across rats (adapted from Leventhal et al., 2012, with permission from Elsevier). **B**, Mean relative beta power of GPe population firing rates in the network model (averaged over 400 simulations), exposed to ramp, sensory and motor stimulation patterns (solid lines). For comparison, if the striatal motor input to GPe is withheld in the model (dashed lines), the second beta peak disappears for long reaction time trials (see blue dashed line in right panel). **C**, Single-trial striatal LFP traces from a single recording session, sorted by reaction time, aligned to the Go cue (left) and movement onset (right) with beta epochs marked in red (adapted from Leventhal et al., 2012, with permission from Elsevier). **D**, Same visualization for single-trial model simulations with each trace showing the population firing rate of GPe neurons in the network model. For simulation of each trial, the model reaction time was randomly selected from the experimental data.

Using our stimulation patterns based on single unit recordings, we studied how different reaction times affect the time course of beta power. We found a strikingly similar effect of reaction time on the time course of beta power in the network model (Figure 6B). For long reaction time trials the model exhibited two separate peaks in the mean beta power with the same time course as the experimental LFP data (Figure 6B). Furthermore, the peak of the mean beta power in the model after movement onset for short reaction time trials had a higher amplitude than in long reaction time trials, similar to the experimental LFP data (see right panels in Figures 6A and 6B). The ability of the model to capture the fine details of the complex beta power modulation became visible even at the single-trial level (Figures 6C, D). As in the experimental data, changes in mean power modulation were reflected as a change in the probability of a transient beta oscillation, rather than as only a gradual increase in the oscillation amplitude.

To understand the mechanisms underlying the complex relationship between beta and reaction times, we can now use our network model to determine the contribution of each stimulation component. Before the Go cue, ramping activity of the STN neurons in the model causes a gradual increase in beta power (mostly because of an increase in the probability of a beta event), starting almost 600 ms before the Go cue (Figures 6B, D and Figure 7). At the time of the Go cue the sensory responses of the STN neurons generate a weak and brief beta oscillation in the model (green traces in Figure 7). In short reaction time trials this brief beta oscillation overlaps with beta oscillations driven by “ramp” and “motor” inputs (as sensory and motor events are temporally close). This overlap results in an interaction of ongoing beta (driven by “ramp” input) with beta driven by “motor” input, leading to high beta power around the time of movement onset (Figures 6B and 7, top). For long reaction time trials, after the Go cue, but before movement initiation, the “sensory” and “ramp” inputs determine the beta dynamics in the model. The interaction between the “sensory” and “ramp” inputs leads to the first, high-amplitude beta peak for long reaction time trials (Figures 6B and 7, bottom). As Go cue and Nose Out events are temporally distant for long trials, this high-amplitude beta power starts to decay before the time of movement onset. This is followed by another beta epoch due to “motor” input which leads to the second peak of beta power, after the time of movement onset, for long reaction time trials (Figures 6B, D and 7). The amplitude of this second peak is smaller, compared to the peak after movement onset for short reaction time trials (Figure 6B, right), because it lacks the interaction with STN excitation due to the Go cue (Figure 7). Functionally, the first beta peak in long reaction time trials may be linked to the prolongation of movement initiation in high beta states (Levy et al., 2002; Brown et al., 2001; Chen et al., 2007; Pogosyan et al., 2009). Thereby our model connects “ramp” activity in STN with the generation of beta oscillations and potential functional roles as a “brake” (Frank, 2006).

**Figure 7:**
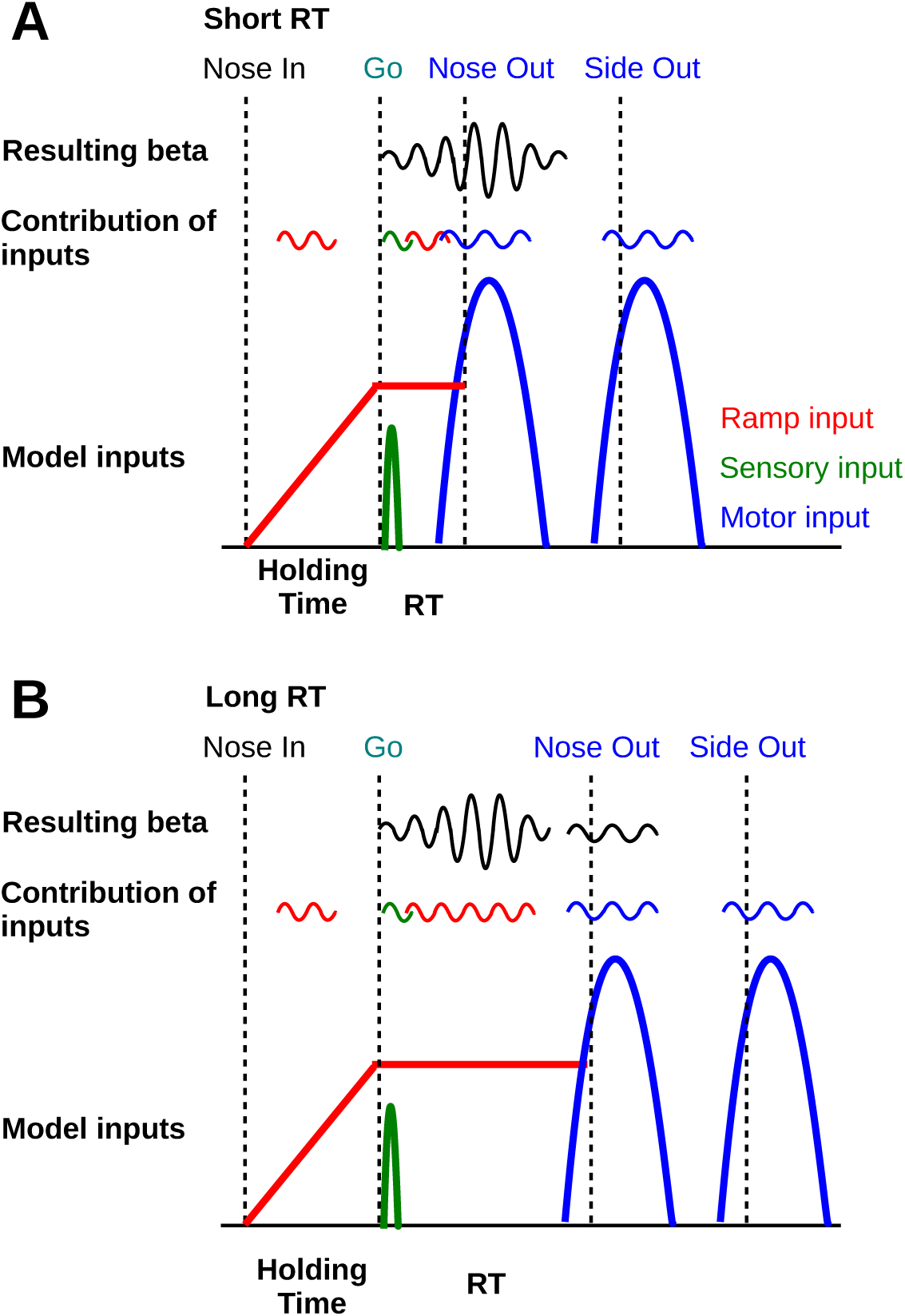
Scheme of contribution of each stimulation component to the generation of beta oscillations in short (A), and long (B) reaction time trials. Red, green, and blue schematized beta oscillations show the contribution of each individual input (ramp, sensory, and motor inputs, respectively) without the other one. Note that for short reaction time trials, interaction between beta oscillations due to ramp, sensory, and motor inputs leads to transient increase in beta power around the time of movement onset (black trace shows the net effect of the interaction). For long reaction time trials, interaction between beta oscillations due to sensory and ramp inputs leads to transient increase in beta power before the time of movement onset which is followed by another beta epoch due to motor input (black traces show the net effect of the interaction).

### Our results are robust to the STN-STN recurrent connectivity in the network model

In the network model we used, the STN neurons received excitatory synaptic inputs from other STN neurons with a connection probability of 2% (Kumar et al., 2011). However, several experimental studies indicate that the STN-STN recurrent connectivity is very rare or do not exist (Hamond and Yelnik, 1983; Sato et al., 2000; Parent and Parent 2007; Koshimizu et al., 2013). Therefore, we modified the network model parameters to test if the model without STN-STN connections is also able to capture the behaviorally relevant dynamics of the LFP beta oscillations. Indeed, with slight modifications of parameters (see Methods), all key results, including the time course of beta around the time of movement preparation and execution (Figure 8A), the beta phase reset (Figures 8B, C), and the complex relationship between beta and reaction time (Figures 8D, E), were reproduced. This demonstrates that our model account of transient beta oscillations does not depend on STN-STN recurrent connectivity.

**Figure 8:**
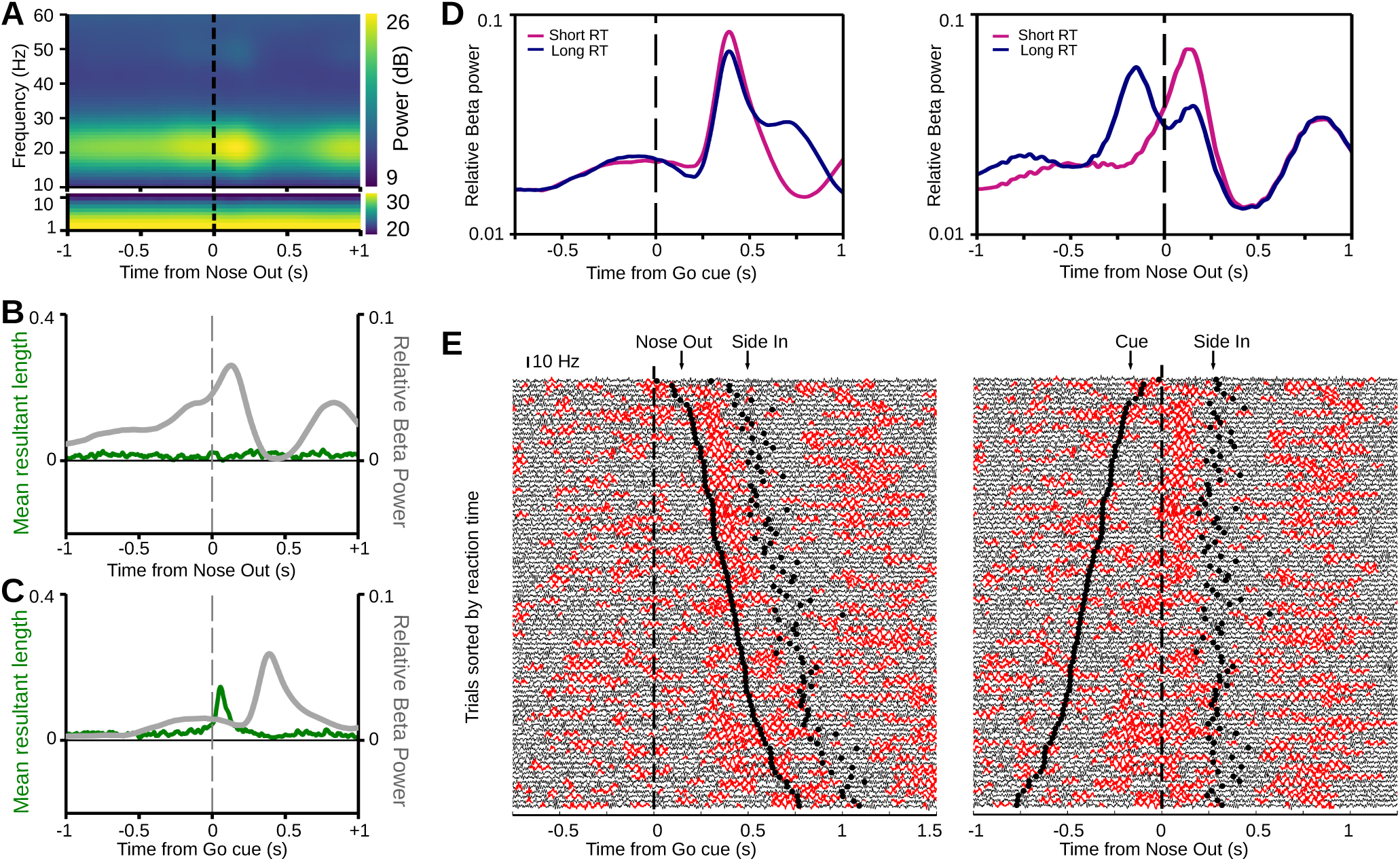
The network model without recurrent connections in STN reproduces all key results. **A**, Mean spectrogram (over 400 simulations) of GPe average firing rates for simulation of correct Go trials in the modified network model matching the time course of beta power in the experimental data. **B, C**, Time resolved beta mean resultant length (left axes, green) and beta power (right axes, gray) of the GPe population firing rate in the modified network model, aligned to the movement onset (B) and to the Go cue (C; average of 400 simulations). **D**, Mean relative beta power of GPe population firing rates in the modified network model aligned to the Go cue (left), and movement onset (right), averaged across 400 simulations. **E**, Single-trial simulations of the modified network model, sorted by reaction time, with each trace showing the population firing rate of GPe neurons, aligned to the Go cue (left) and movement onset (right; beta epochs are marked in red).

In summary, our results show that the combination of 1) sensory responses of STN neurons, 2) movement-related inhibition of GPe neurons, and 3) ramping activity in STN, account for the complex properties of beta power modulation over time, beta phase reset and correlations with reaction time of rat electrophysiological recordings in the basal ganglia. Thereby, the model allows us to make clear predictions about the underlying mechanisms and provides the basis for studying functional consequences on neural processing and behavior.

## Discussion

Oscillations in the LFP often reflect sensory, cognitive and motor aspects of neural processing, but we lack understanding of how and why network oscillations emerge. Furthermore, we face a gap between firing patterns of single neurons and network dynamics. Here we addressed this by combining experimental data with computational modeling to study how firing patterns in single units of task-performing healthy rats affect basal ganglia network dynamics. Although our computational model was originally used to describe beta oscillations in Parkinson’s disease, this model also accounted for properties of beta in healthy animals. Thereby, we characterize potential neuronal mechanisms underlying oscillations, relate healthy to pathological beta oscillations, and provide avenues for studying functional roles of beta in behavior.

### Neuronal mechanisms of beta oscillations

Computational and experimental studies have implicated the STN-GPe network in beta oscillations in Parkinson’s disease (Brown et al., 2001; Magill et al., 2001; Terman et al., 2002; Bevan et al., 2002; Rubin and Terman, 2004; Brown and Williams, 2005; Mallet et al., 2008a; Tachibana et al., 2011; Stein and Bar-Gad, 2013; Nevado-Holgado et al., 2014; Pavlides et al., 2015; Wei at al., 2015). Moreover, cortico-subthalamic excitation as well as striato-pallidal inhibition can generate beta oscillations in network models of the subthalamo-pallidal loop (Gillis et al., 2002; Kumar et al., 2011; Nevado-Holgado et al., 2014; Pavlides et al., 2015; Wei at al., 2015; Ahn at al., 2016). Consistently, we show that temporally regulated subthalamic excitation and pallidal inhibition reproduces the dynamics of transient beta oscillations observed in the healthy basal ganglia during behavior. Therefore, the same network that is responsible for beta oscillations in Parkinson’s disease may also be involved in the generation of healthy beta.

As an alternative to the STN-GPe network, striatal MSNs (McCarthy et al., 2011), feedback projections from GPe back to striatum (Corbit et al., 2016), or spread of cortical beta to STN may be involved in basal ganglia beta oscillations. However, our model supports the role of the STN-GPe network due to the close correspondence between single unit activity and the resulting complex time course of beta oscillations. Whether other models for the generation of beta would be able to account for the complex time course and behavioral correlates of beta remains to be shown. While increased striatal spiking increases beta oscillations in several models (McCarthy et al., 2011; Kumar et al., 2011; Corbit et al., 2016), our model emphasizes the role of excitatory inputs to STN for the transient dynamics of beta oscillations. Overall, as beta oscillations are a heterogeneous phenomenon (Szurhaj et al., 2003; Kilavik et al., 2012; Feingold et al., 2015), cortical and subcortical circuit may contain several mechanisms for the generation of beta, e.g. to permit long range communication (Fries, 2005). Therefore, these models are not necessarily exclusive and a key future challenge will be to disentangle the different circuits and their interaction. Nonetheless we have shown that the STN-GPe network is sufficient to explain many features of beta oscillations in awake behaving animals.

### Direct and indirect pathway MSNs

Activity of direct pathway MSNs (striato-nigral) promote actions, while indirect pathway MSNs (striato-pallidal) suppress actions (Albin et al., 1989; Alexander and Crutcher, 1990; Kravitz et al., 2010; Freeze et al., 2013; Roseberry et al., 2016). Here we considered movement-related increases in MSN activity (Figure 1E) as inhibitory input to the model GPe (Figures 3A, B), without knowing whether the recorded MSNs are part of the direct or indirect pathway. This assumption is supported by evidence that direct and indirect pathway MSNs are concomitantly active during movements (Cui et al., 2013; Isomura et al., 2013). Nevertheless, there might be important activity differences between direct and indirect pathway neurons coordinating behavior. Whether co-activation of indirect pathway MSNs during movement reflects the suppression of alternative actions (Hikosaka et al., 2006; Redgrave et al., 2010) or activates specific neural assemblies in motor cortex (Oldenburg and Sabatini, 2015) remains unclear. Furthermore, almost 60% of direct pathway MSNs, possess collateral terminal fields in GPe (Cazorla et al., 2014). Therefore, during movements GPe likely receives increased inhibitory input from striatal MSNs as incorporated in the model.

### STN as a brake

We found ramps in the activity of STN units while the animal was waiting for the Go cue. During this time the animal has to prevent premature movements to receive the food reward. Building on “hold-your-horses” models of STN (Frank, 2006), these ramps might prevent or delay movements. Correspondingly, in our experimental data the ramps reached a plateau after the Go cue, which was linked to the reaction time (i.e. the plateau persisted longer in trials with a long reaction time; Figure 2A, bottom). Therefore, these ramps might modulate the readiness for movement initiation. However, we also observed (data not shown) that the population activity of the STN ramps did typically last until movement initiation, indicating that the offset of this STN ramp does not provide a motor command itself. Instead, high STN activity might ensure that only coordinated movement commands (potentially signaled by striatal output), but not premature movement impulses, lead to motor output.

Conceptually, our model provides an important link between putative “hold-your-horses” ramping activity in STN, beta oscillations and reaction times. The ramping activity increased spiking activity of the STN neurons and, consequently, lead to also more beta oscillations in the model (Kumar et al., 2011). This was key in accounting for the bimodal shape of the mean beta power for long reaction time trials (Figure 6B).

The STN ramps might be due to cortical drive. For example, in the motor cortex of monkeys ramping activity has been observed while the animals anticipated sensory cues and needed to prevent premature movements (Confais et al., 2012). Furthermore, other cortical areas including right inferior frontal cortex and the pre-supplemental motor area project to STN and have been implicated in motor suppression (Wessel and Aron, 2017). In general, cortico-subthalamic excitation has previously been proposed to be important for the generation of beta oscillations (Tachibana et al., 2011; Pavlides et al., 2015). Importantly, the STN ramps during the hold period increased the probability of transient beta in our model. This fits well with anti-kinetic aspects of beta (Brown and Williams, 2005), and with STN activity correlating with slowness of movement observed during the progression of Parkinson’s disease (Bergman et al., 1994; Remple et al., 2011).

### Behavioral relevance and predictions

Beta oscillations seem to comprise a heterogeneous phenomenon with potentially different functions and mechanisms depending on the brain region (Szurhaj et al., 2003; Kilavik et al., 2011; Feingold et al., 2015). Here we extend this view by proposing that transient, non-pathological basal ganglia beta can be driven by two distinct inputs. Firstly, beta oscillations were driven by excitatory inputs to STN, including the ramping activity that might be linked to preventing premature movements. Secondly, beta oscillations were also driven by striato-pallidal inhibition during movement. Therefore, our model provides an explanation for why beta in some cases can be “antikinetic” (Brown and Williams, 2005), but in other cases can also appear during movement (Leventhal et al., 2012). Whether and how these two modes of beta make different functional contributions, e.g. by differential communication with other brain regions (Fries, 2005), is an open question.

Based on our model we make several experimentally testable predictions. Firstly, the two modes of beta generation, via GPe inhibition and STN excitation, might have different signatures in LFP recordings. If the beta is generated by GPe inhibition, the oscillation begins with a decrease in GPe activity. If beta is generated by STN excitation, the beta oscillation begins with an increase in STN. Although we do not know yet how spiking in the STN and GPe relates to patterns in the LFP, these two modes could translate into different onset phases of beta. Therefore, we presume that transient beta oscillations could be classified based on their onset phase, and that this is indicative of whether the oscillation was driven by input to GPe or STN. Despite practical challenges, such as detecting the exact onset phases of beta in noisy LFPs, this might provide valuable insights into whether the two modes of beta generation have distinct behavioral correlates.

Secondly, our model makes specific predictions about the relation between activity of MSNs projecting to GPe and the timing of beta oscillations (McCarthy et al., 2011). In recordings of identified direct and indirect pathway MSNs, our model predicts that the activity of the D2 MSNs predicts the timing of beta more accurately than the activity of the D1 MSNs. One complicating factor is that this distinction does not apply to beta driven by cortical excitation of STN.

Another model prediction arises from our observation that the duration of excitatory inputs to STN determines whether a phase reset occurs in the LFP or not. Sensory neuronal responses (Figures 1C, D) are typically brief. We propose that sensory cues from other modalities have the same effect, so that e.g. visual cues that lead to brief excitations of STN also lead to a phase reset in the LFP signal. Furthermore, in addition to sensory cues, brief optogenetic stimulation of STN might yield the same effect. Whether these cue-induced beta phase resets play also a functional role, e.g. in the temporal coordination with inputs from other regions, remains to be shown.

Finally, we predict that changes in the structure of the STN ramping activity affects the probability of beta oscillations. If the STN ramps indeed reflect a “hold your horses” signal (Frank, 2006), changes in the behavioral paradigm that manipulate the readiness for movement initiation should directly affect the ramping activity. For example, if the cost for the animal of a premature response is increased, the corresponding ramping activity might change its time course and amplitude. In the model this would directly translate into changes in the time course and probability of transient beta.

In conclusion, the direct combination of our computational model with experimental data provides a connection between single unit activity and network oscillations. This helps us to study the functional contributions of transient beta oscillation during sensorimotor processing in a behavioral context.

## Acknowledgments

We thank Wei Wei, Alejandro Jimenez, Lars Hunger, and Mohammad Mohagheghi Nejad for useful comments and discussion. This work was supported by BrainLinks-BrainTools Cluster of Excellence funded by the German Research Foundation (DFG, grant number: EXC 1086) and the University of Sheffield. We also acknowledge support by the state of Baden-Wxürttemberg through bwHPC and the German Research Foundation (DFG) through grant no INST 39/963-1 FUGG.

